# Network pharmacology, single gene survival analysis and molecular docking to study the mechanism of Sotetsuflavone in the treatment of pancreatic cancer

**DOI:** 10.1101/2024.04.30.580419

**Authors:** Zi-Yong Chu, Xue-Jiao Zi

**Author notes:** Corresponding author: Zi-Yong Chu, College of Life Science and Technology, Xinjiang University, Urumqi,830046, Xinjiang, PR China.

## Abstract

Pancreatic cancer is a highly lethal cancer with limited treatment options. The number of pancreatic cancer patients is increasing rapidly worldwide. Many natural products have been shown to have anticancer activity in a range of studies. Sotetsuflavone is derived from *Cycas revoluta* Thunb. and exhibits anticancer activity. The present study incorporates network pharmacology, single gene survival analysis, gene expression analysis and molecular docking to reveal the mechanism of Sotetsuflavone in the treatment of pancreatic cancer. Initially, it was evaluated through multiple databases for a comprehensive pharmacological evaluation of Sotetsuflavone. Then, the target information of Sotetsuflavone and pancreatic cancer was searched and screened using public databases. According to the results of matching the potential targets of Sotetsuflavone with the targets of pancreatic cancer, the protein-protein interaction (PPI) network was constructed by using the STRING database and imported into Cytoscape 3.9.0 for the network analysis to screen the hub targets, and then classify and co-expression analysis was carried out on the hub targets. Then, we conducted Gene Ontology (GO) analysis and Kyoto Encyclopedia of Genes and Genomes (KEGG) pathway enrichment analyses of potential and hub targets and discovered that these targets were involved in pancreatic cancer-related signalling pathways. In addition, the GEPIA 2 platform was used to be performed for single gene survival analysis and gene expression analysis, and AutoDock Vina was employed for molecular docking analysis. In total, we have acquired 31 hub targets for the treatment of pancreatic cancer by Sotetsuflavone, namely ABCB1, AURKA, CDK1, and so on. Kaplan-Meier survival analyses demonstrated that ABCB1, AURKA, CDK1, HDAC6, MET, and MMP3 are promising hub targets that can be used as biomarkers for pancreatic cancer diagnosis and prognosis. These hub targets are highly expressed in pancreatic cancer tissues compared to normal tissues. The molecular docking results showed a strong binding capacity of Sotetsuflavone to these hub targets. In summary, it is proposed that Sotetsuflavone is a new anticancer drug, which can regulate cancer-related signalling pathways such as pancreatic cancer by inhibiting the activities of ABCB1, AURKA, CDK1, HDAC6, MET, and MMP3, which are hub targets with up-regulated expression in pancreatic cancer tissues, in order to treat pancreatic cancer. However, it also requires a series of in vivo and in vitro studies to ensure its safety and efficacy.

## 1. Introduction

Pancreatic cancer is the seventh cause of cancer deaths globally, with more than 430,000 people died of pancreatic cancer each year[1]. There has been more than a doubling of the number of pancreatic cancer patients worldwide in the last 25 years[2]. It has recently been demonstrated that abnormal microbiological metabolism in humans, blood type, blood glucose and lipid levels are significant factors in the occurrence of pancreatic cancer[3–5]. The factors such as area, gender, age and genetics are also associated with the propagation of this disease[6–9]. The only current hope for curing pancreatic cancer, with its extreme aggressiveness and poor prognosis, is surgery[10]. There are limited treatment alternatives and high death rates for pancreatic cancer that have attracted a significant amount of attention from researchers all around the world. Therefore, the discovery of new anticancer drugs is immediately urgent.

Sotetsuflavone is a flavonoid compound isolated from *Cycas revoluta* Thunb.[11]. According to reports, Sotetsuflavone has antibacterial, anticancer, anti-inflammatory and antiviral effects[12–14]. It is noteworthy that Sotetsuflavone can inhibit the invasion and metastasis of non-small cell lung cancer A549 cells by reversing EMT through TNF-α/NF-κB and PI3K/AKT signalling pathways[15], and also enhances the inhibitory effect of 5-fluorouracil on gastric cancer cells through the AKT-mTOR pathway[16]. Therefore, it is suggested that Sotetsuflavone is a potential anticancer drug and is promising to be a new drug for the treatment of pancreatic cancer.

Network pharmacology is a novel approach to the systematic research of drugs that enables the study of drug interactions with multiple targets and the clarification of their mechanisms of action[17]. The fundamental idea of network pharmacology is to utilise big data information to analyse the treatment effects of drugs on diseases on the basis of “Drug-Target-Pathway”[18]. There is a variety of applications of current network pharmacology for diseases such as inflammation, cancer, cardiovascular disease, neurodegenerative disease, and so on[19]. For instance, Shreni Agrawal et al.[20] used network pharmacology to study the therapeutic effects of kaempferol and catechins on pancreatic cancer.

Network pharmacology was used in this research to establish a multi-target, multi-pathway model to explain the interactions between Sotetsuflavone and pancreatic cancer targets from a network viewpoint. Single gene survival analyses, gene expression analyses and molecular docking studies were supplemented with validation of the results. In the present study, for the first time, a comprehensive evaluation of Sotetsuflavone was carried out using several pharmacological databases to support further research on the pharmacological effects of Sotetsuflavone. In addition, it was the first time that we explored the mechanism of action of Sotetsuflavone in the treatment of pancreatic cancer, providing theoretical support and direction for further basic research.

## 2. Methods

### 2.1. Pharmacological evaluation of Sotetsuflavone

For pharmacological evaluation of Sotetsuflavone, its standard structure and SMILE nodes were obtained from PubChem database. Then the structural formulae of Sotetsuflavone was imported into SwissADME database(http://www.swissadme.ch/)[21], ADMETlab 2.0 database(https://admetmesh.scbdd.com/)[22], Tox-Prediction-II database(https://tox-new.charite.de/protox_II/index.php?site=compound_input)[23], and Molinspiration database(https://molinspiration.com/)[24] for Physicochemical Properties, Pharmacokinetics, Pharmacotoxicity, and Bioactivity analyses, respectively.

### 2.2. Targets for collection of Sotetsuflavone

The standard structure and SMILE nodes of Sotetsuflavone were determined by searching for “Sotetsuflavone” using the PubChem database. Based on the search results, we imported the structural information of Sotetsuflavone into the SwissTargetPrediction database(http://swisstargetprediction.ch/)[25] and selected the species “Homo sapiens” to predict the potential targets. In addition, we also imported the structural information of Sotetsuflavone into the TargetNet database(http://targetnet.scbdd.com/home/index/)[26] and selected “AUC ≥0.7” and “Fingerprint type: ECFP4 Fingerprints” to predict the potential targets. The predictions from the two databases were then imported into the Uniport database(https://www.uniprot.org/)[27] for standardisation before merging the results and removing duplicate potential targets. Finally, these merged targets were used to construct a target library for Sotetsuflavone.

### 2.3. Screening for pancreatic cancer-related targets

Targets related to pancreatic cancer were collected from the GeneCards database (https://www.genecards.org)[28] using “pancreatic cancer” as keyword. Additional targets related to pancreatic cancer were collected from DisGeNET database (https://www.disgenet.org)[29].The predictions from both databases were then imported into the Uniport database(https://www.uniprot.org/)[27] for normalisation before merging and removing duplicate targets. Then, the results were used to construct a library of targets for pancreatic cancer.

### 2.4. Construction of protein interaction network and screening of hub targets

Firstly, we used the Vene diagram to screen the common potential targets of Sotetsuflavone and pancreatic cancer, and regarded the intersection part as a potential target of Sotetsuflavone for the treatment of pancreatic cancer. Then, the intersecting targets were imported into STRING database(https://cn.string-db.org/)[30] and the species “Homo sapiens” was selected to construct the protein-protein interaction (PPI) network and the results were imported into Cytoscape 3.9.0[31] for network analysis.

We used the following criteria to screen the hub targets of Sotetsuflavone for the treatment of pancreatic cancer: ① Betweenness Centrality > median, ② Closeness Centrality > median, ③ Degree ≥ twice median ④ Radiality > median, and ⑤ Topological Coefficient > median[32]. The hub targets were then imported into the STRING database(https://cn.string-db.org)[30] for network analysis and co-expression analysis.

### 2.5. Functional annotation and pathway analysis

In order to investigate the biological function of potential targets of Sotetsuflavone for the treatment of pancreatic cancer, the DAVID database(https://david.ncifcrf.gov/summary.jsp)[33] was used for Gene Ontology (GO) analysis and Kyoto Encyclopedia of Genes and Genomes (KEGG) pathway enrichment analysis[34]. We elucidated major biological functions through Gene Ontology (GO) analyses, including the assessment of biological processes (BP), cellular components (CC) and molecular functions (MF)[34]. In addition, we investigated the signalling pathways related to the treatment of pancreatic cancer with Sotetsuflavone by KEGG enrichment analysis[35].

Meanwhile, we also performed KEGG enrichment analysis of the hub targets of Sotetsuflavone for the treatment of pancreatic cancer using the DAVID database. This analysis was aimed at further investigating the pathways involved in the hub targets of Sotetsuflavone for the treatment of pancreatic cancer in order to elucidate and highlight the important signalling pathways involved. Finally, we visualised the results of the GO and KEGG enrichment analyses.

### 2.6. Survival analysis

We have conducted survival analyses of the hub genes using GEPIA 2 (http://gepia2.cancer-pku.cn/)[36]. The clinical significance of a particular gene can also be assessed by survival analyses based on gene expression levels[19]. In the GEPIA 2 server, the Kaplan-Meier survival curves were constructed using a database of pancreatic cancer cells to examine the relationship between hub targets and overall survival of patients with pancreatic cancer. The gene normalisation function available in GEPIA 2 allows the relative expression of two different genes as input[19]. We considered Hub targets with p-value < 0.01 to be statistically significant and performed expression analysis and molecular docking analysis.

### 2.7. Gene expression analyses

To investigate the expression of hub targets in normal and pancreatic cancer tissues, we used the GEPIA 2 platform(http://gepia2.cancer-pku.cn/) to analyse the differences in the expression of hub targets with p-value < 0.01 at the RNA level in normal and pancreatic cancer tissues[36].

### 2.8. Molecular docking analysis

The molecular docking approach was used to validate the existence of interactions between Sotetsuflavone and hub targets with p-value < 0.01 for therapeutic efficacy in pancreatic cancer. First, we converted the previously acquired structure files of Sotetsuflavone from sdf format to pdb format using ChemBioDraw 3D software[37]. Then, we used AutoDock Tools 1.5.7 to pre-process the molecular docking of the hub targets and Sotetsuflavone downloaded from the RCSB Protein Data Bank (PDB)(https://www.rcsb.org/)[38]. Next, we used the ProteinsPlus platform(https://proteins.plus/)[39, 40] to predict the size and location of the molecular docking activity pocket of the hub targets. We then ran AutoDock Vina for molecular docking using the Anaconda Powershell Prompt command prompt[41]. Finally, we visualised and analysed the results using Discovery studio visualizer, Maestro 12.8 and PyMOL and plotted the binding energy heatmap of molecular docking.

### 2.9. Statistical analysis

There were data for the synthesis study from an online database platform and visualised and analysed using the SRplot online mapping platform[42].

## 3. Results

### 3.1. Pharmacological assessment of Sotetsuflavone

The SwissADME database(http://www.swissadme.ch/), ADMETlab 2.0 database(https://admetmesh.scbdd.com/), Tox-Prediction-II database(https://tox-new.charite.de/protox_II/index.php?site=compound_input) and Molinspiration database(https://molinspiration.com/) were utilised for the pharmacological evaluation of Sotetsuflavone (**Fig. 2**). Drug-likeness prediction usually requires consideration of the following criteria, such as optimised Lipinski Rule of 5[43]. The Physicochemical Properties analyses we carried out on Sotetsuflavone using SwissADME database showed that the Molecular weight (MW) of Sotetsuflavone < 600, Number of hydrogen bond acceptors < 12, Number of hydrogen bond donors < 7. The pharmacokinetic prediction of Sotetsuflavone using ADMETlab 2.0 database showed that Caco-2 Permeability < -5.15 Log unit, Bioavailability < 30%, Blood-Brain Barrier Penetration = 0.001, Plasma Protein Binding > 90%, T_1/2_ >3h, and Clearance < 5 mL/min/kg. The Tox-Prediction-II database was used to predict the toxicity of Sotetsuflavone. The results showed that Sotetsuflavone had activated toxicity such as Immunotoxicity, Aryl hydrocarbon Receptor, and Mitochondrial Membrane Potential. However, Hepatotoxicity, Mutagenicity and Cytotoxicity were not activated. Although Sotetsuflavone are toxic, an LD50 of 4000 mg/kg is required to be effective. The Molinspiration database was used to analyse the bioactivity of Sotetsuflavone. When the bioactivity score is greater than 0, between -0.5 and 0, or less than -0.5, it is considered active, moderately active, or inactive, respectively[44]. The bioactivity scores of Sotetsuflavone showed activity except Ion channel modulator which showed moderate activity. These results indicated that Sotetsuflavone is a potential drug and can be safely used in biotherapeutics (**Fig. 2**).

**Fig. 1.**
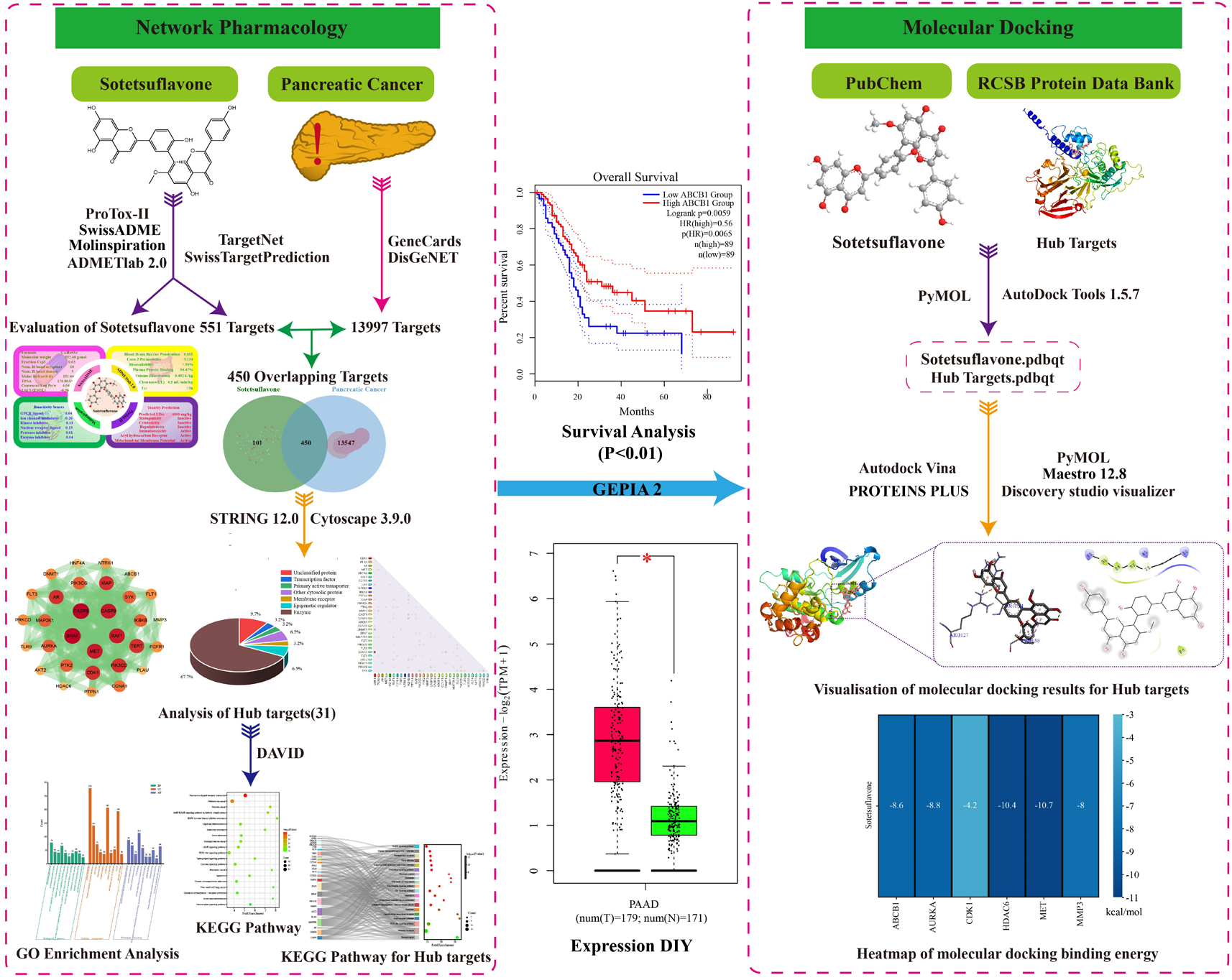
General workflow of the study.

**Fig. 2.**
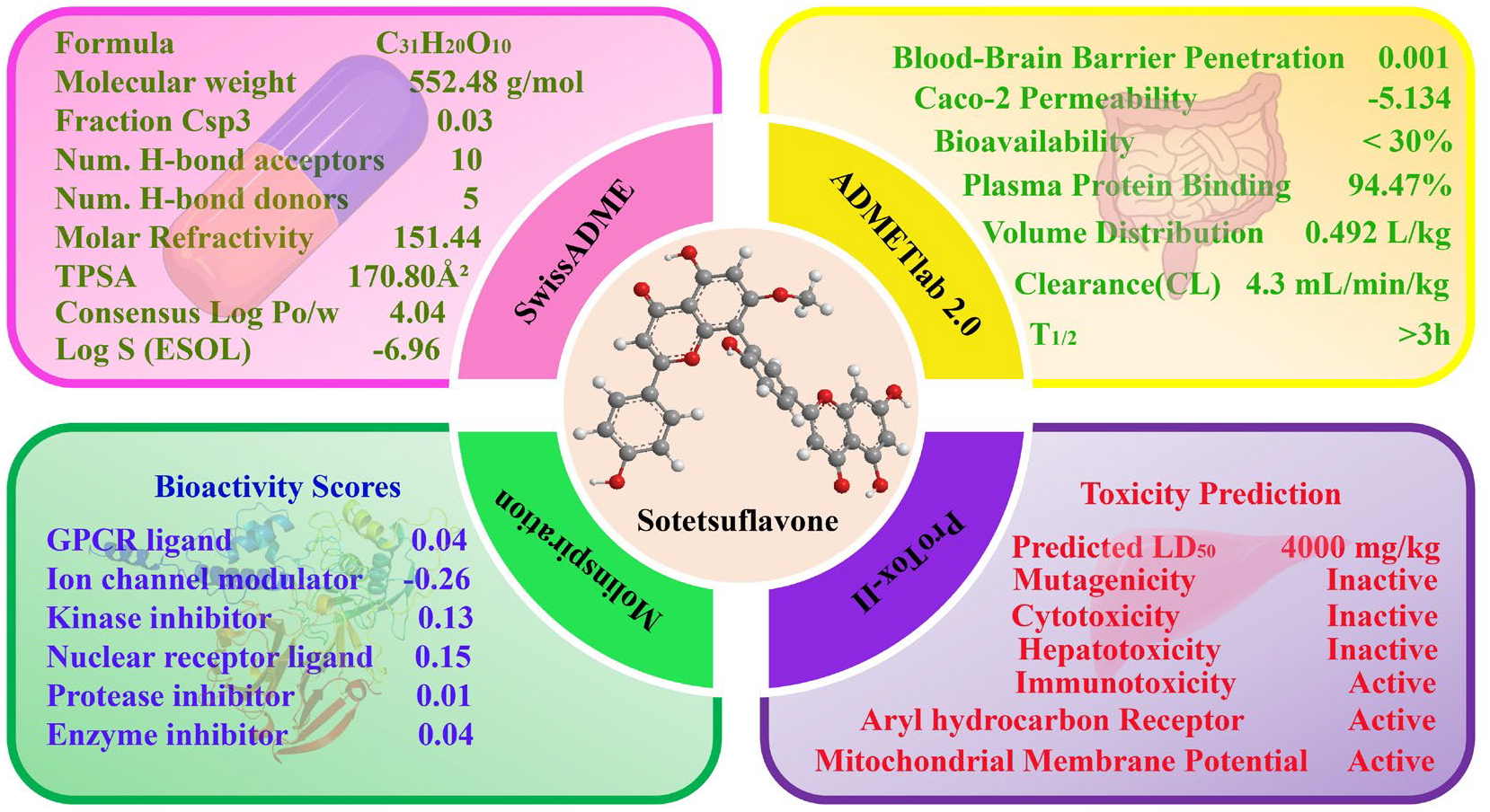
The pharmacological evaluation of Sotetsuflavone.

### 3.2. The prediction and screening of targets

The targets of Sotetsuflavone were predicted and screened using SwissTargetPrediction database(http://swisstargetprediction.ch/) and TargetNet database (http://targetnet.scbdd.com/home/index/), and a total of 551 potential targets were screened. After that, there were 13997 pancreatic cancer-related targets retrieved and screened using GeneCards database (https://www.genecards.org) and DisGeNET database (https://www.disgenet.org). The potential targets of Sotetsuflavone and related targets of pancreatic cancer were finally submitted to the Venn diagram to identify overlapping targets. Overall, a total of 450 overlapping targets were eventually screened, as shown in **Fig. 3**.

**Fig. 3.**
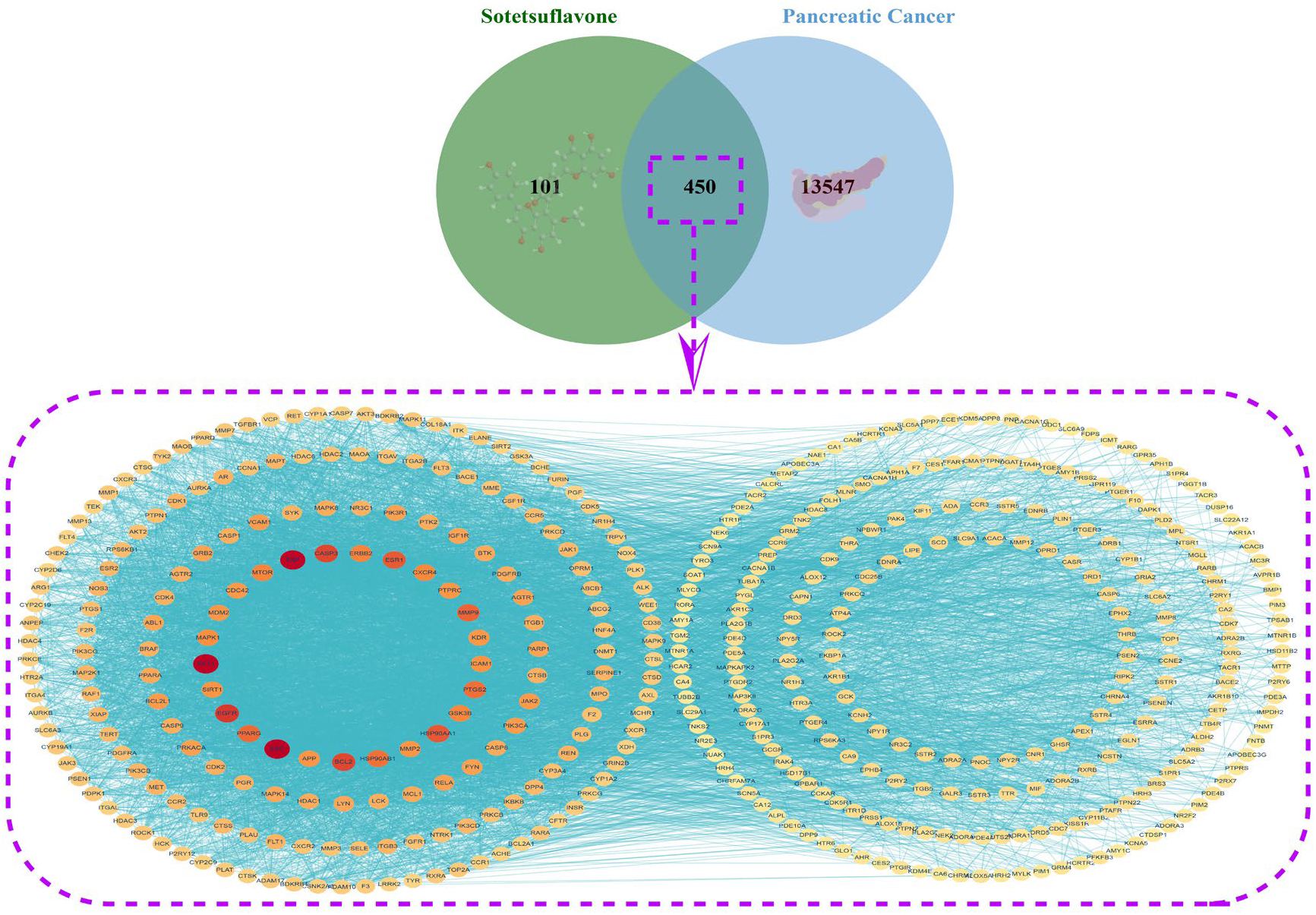
Venn diagram and PPI network of the targets of Sotetsuflavone and pancreatic cancer.

### 3.3. The targets interaction network and the acquisition of hub targets

To construct the PPI network, 450 overlapping targets were searched in the STRING database(https://cn.string-db.org/). The network was used to demonstrate the interactions between targets during disease progression. Subsequently, there were 31 hub targets screened by importing the PPI network files into Cytoscape 3.9.0 for network analysis (**Table 1**, **Fig. 4A**, **Fig. 4B**). Then, we categorized the 31 hub targets and found that they belonged to the highest number of enzyme targets, followed by unclassified proteins, Epigenetic regulator, Other cytosolic protein, etc., which accounted for 67.7%, 9.7%, 6.5%, and 6.5%, respectively (**Fig. 4C**). Furthermore, Co-expression analyses of the hub targets indicated co-expression between these targets (**Fig. 4D**). Through the analysis of connectivity patterns and interactions in this network, it provided insights into the molecular interactions involved in the treatment of pancreatic cancer with Sotetsuflavone.

**Fig. 4.**
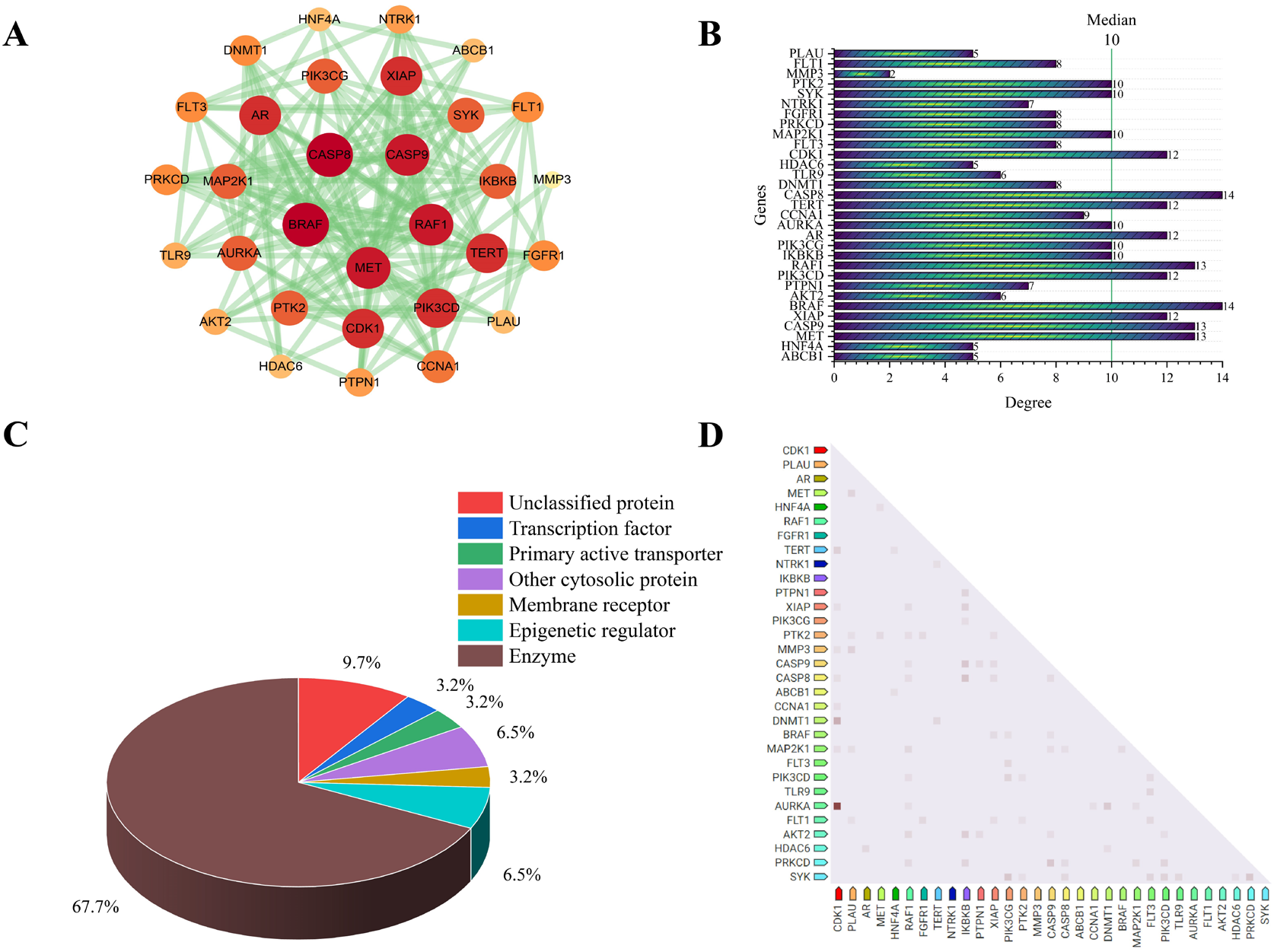
Multi-target analysis of network pharmacology-based Sotetsuflavone for the treatment of pancreatic cancer. (A) The PPI network of hub targets. (B) The PPI network bar plot of hub targets. (C) The classification of hub targets. (D) The co-expression of hub targets in Homo sapiens.

**Table 1.** Hub targets screened from PPI network.

| Targets | Betweenness<br>Centrality | Closeness<br>Centrality | Degree | Radiality | Topological<br>Coefficient |
| --- | --- | --- | --- | --- | --- |
| CASP8 | 0.002933 | 0.520979 | 75 | 0.995782 | 0.199446 |
| BRAF | 0.004018 | 0.519164 | 68 | 0.995751 | 0.197438 |
| CASP9 | 0.002478 | 0.518561 | 71 | 0.995741 | 0.202948 |
| SYK | 0.001705 | 0.518561 | 67 | 0.995741 | 0.194391 |
| FLT1 | 0.001555 | 0.516763 | 65 | 0.995710 | 0.195828 |
| MET | 0.002316 | 0.516166 | 65 | 0.995700 | 0.202144 |
| AR | 0.002916 | 0.514977 | 66 | 0.995680 | 0.193858 |
| PTK2 | 0.002217 | 0.513793 | 73 | 0.995659 | 0.197583 |
| MAP2K1 | 0.001237 | 0.510857 | 63 | 0.995608 | 0.211810 |
| IKBKB | 0.003398 | 0.510274 | 56 | 0.995598 | 0.199564 |
| PIK3CG | 0.001988 | 0.509692 | 57 | 0.995587 | 0.198945 |
| PTPN1 | 0.002221 | 0.507378 | 52 | 0.995546 | 0.193015 |
| ABCB1 | 0.003828 | 0.505656 | 51 | 0.995515 | 0.196465 |
| XIAP | 0.001820 | 0.505085 | 61 | 0.995505 | 0.206533 |
| PLAU | 0.001429 | 0.505085 | 52 | 0.995505 | 0.200570 |
| CDK1 | 0.002791 | 0.502247 | 64 | 0.995454 | 0.200434 |
| HNF4A | 0.002842 | 0.502247 | 49 | 0.995454 | 0.192485 |
| AKT2 | 0.000760 | 0.501121 | 51 | 0.995433 | 0.220234 |
| PIK3CD | 0.001580 | 0.500560 | 60 | 0.995423 | 0.193745 |
| TERT | 0.001295 | 0.500000 | 54 | 0.995413 | 0.215125 |
| NTRK1 | 0.003267 | 0.500000 | 49 | 0.995413 | 0.218250 |
| TLR9 | 0.000844 | 0.498884 | 47 | 0.995392 | 0.201064 |
| RAF1 | 0.000909 | 0.497219 | 57 | 0.995362 | 0.215435 |
| DNMT1 | 0.001064 | 0.494469 | 47 | 0.995310 | 0.217391 |
| PRKCD | 0.001197 | 0.493377 | 49 | 0.995290 | 0.202927 |
| FLT3 | 0.000968 | 0.491749 | 52 | 0.995259 | 0.222765 |
| FGFR1 | 0.001959 | 0.491749 | 49 | 0.995259 | 0.211469 |
| MMP3 | 0.001966 | 0.490670 | 47 | 0.995238 | 0.194767 |
| CCNA1 | 0.001746 | 0.487459 | 50 | 0.995177 | 0.200160 |
| AURKA | 0.001201 | 0.468553 | 47 | 0.994797 | 0.218738 |
| HDAC6 | 0.001012 | 0.490670 | 46 | 0.995238 | 0.219033 |

### 3.4. Functional and pathway enrichment analysis of targets

We conducted GO analyses of 450 potentially overlapping targets using the DAVID database(https://david.ncifcrf.gov/summary.jsp), restricting the species to “Homo sapiens”. With a total of 1547 statistically significant GO entries, there were 1107 biological processes (BPs), 154 cellular components (CCs), and 286 molecular functions (MFs) in our analyses, respectively. The ranking of GO items based on the false discovery rate (FDR) values, and the top 10 items with the lowest FDR values in BP, CC, and MF were selected and visually depicted in the enrichment analysis diagram (**Fig. 5**). Besides, we also conducted KEGG analysis of these 450 potential overlapping targets using the DAVID database to identify their involvement in specific signalling pathways. Within the total of 177 signalling pathways enriched, we generated significance statistical bubble diagrams and categorical histograms (**Fig. 6**), as well as the top 20 KEGG signalling pathways with the lowest FDR values.

**Fig. 5.**
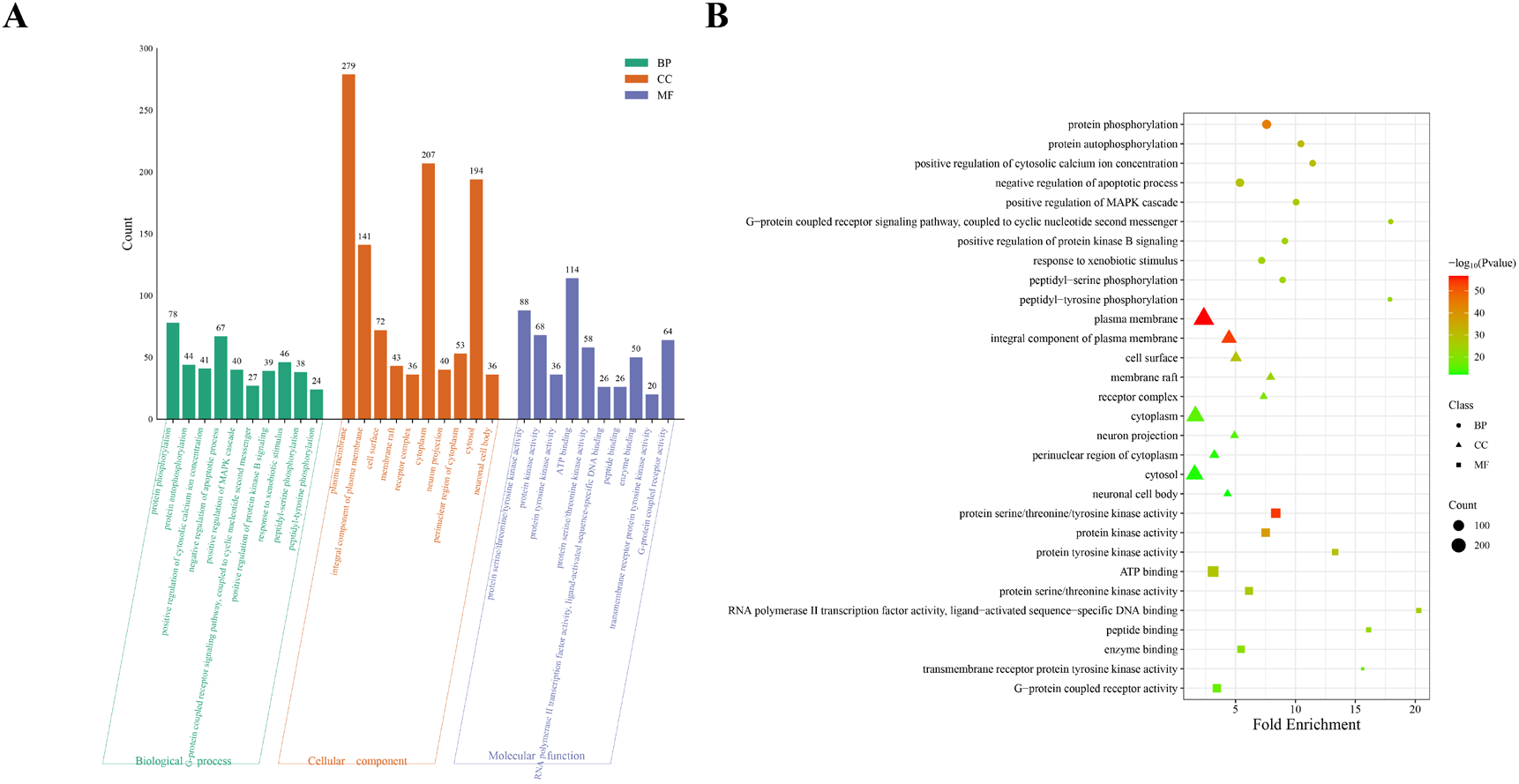
GO enrichment analysis of potential targets (top 10). (A) This histogram illustrated the top 10 enriched entries for each GO category (BP, CC, and MF) with smaller FDR values on the 450 targets. (B) The size of each bubble corresponded gene expressions in a particular pathway.

**Fig. 6.**
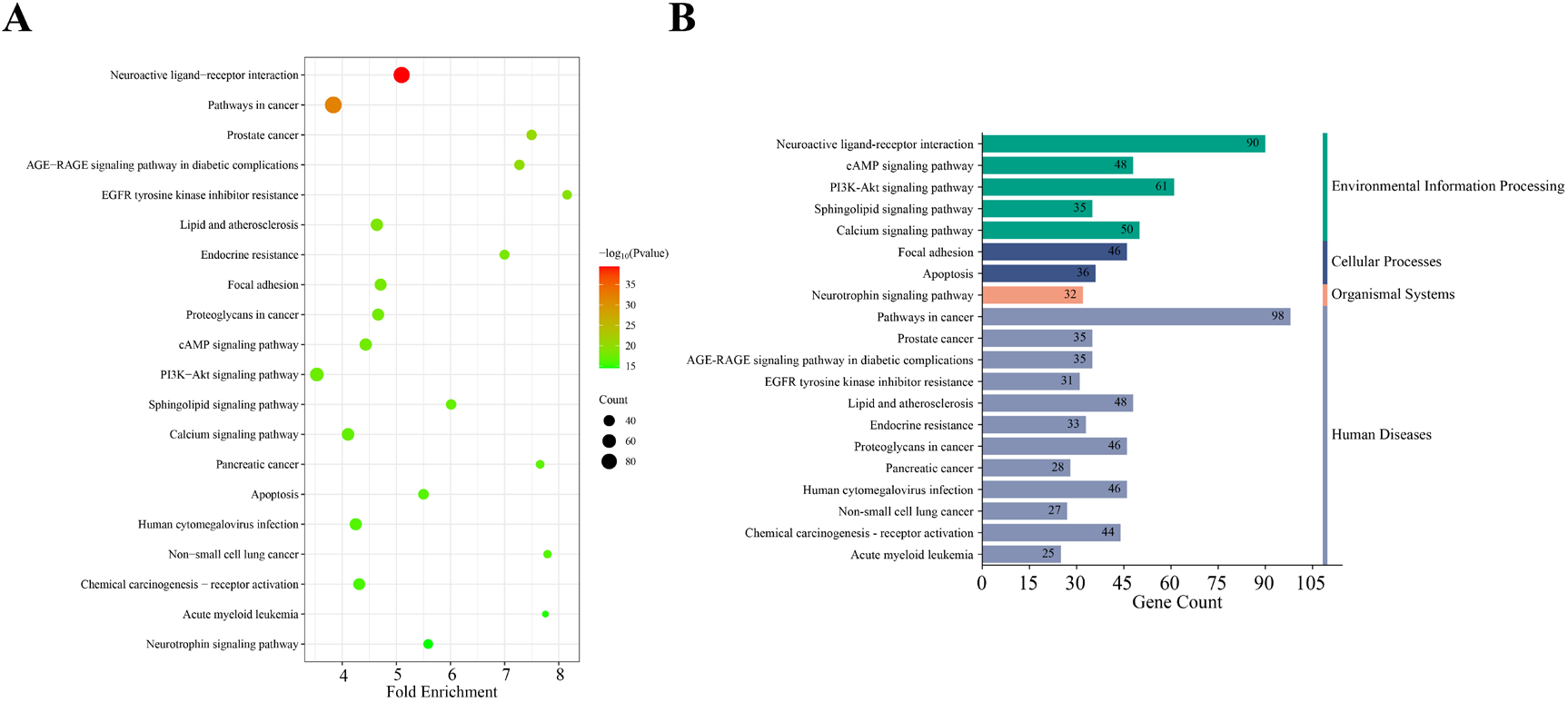
KEGG enrichment analysis of potential targets (top 20). (A) The bubble diagram visualized the top 20 enriched KEGG signal pathways in reverse order of FDR values on the 450 targets. (B) The histogram illustrated the frequency and significance of enrichment for each pathway.

The results obtained from GO and KEGG analyses of potential overlapping targets, based on the results, it is noteworthy that these genes showed a wide distribution and expression in various subcellular localisations and exerted a variety of biological functions, such as G-protein coupled receptor activity, ATP binding, RNA polymerase II transcription factor activity, etc. Moreover, many of these genes are involved in key regulatory processes, such as protein phosphorylation, positive regulation of cytosolic calcium ion concentration and signal transduction, etc. The KEGG signalling pathway enrichment has revealed signalling pathways associated with a variety of signalling pathways, including Neuroactive ligand-receptor interaction, Pathways in cancer, AGE-RAGE signalling pathway in diabetic complications, and Pancreatic cancer. The above results demonstrated that Sotetsuflavone plays an important role in the treatment of pancreatic cancer.

### 3.5. Analysis of pathway enrichment for hub targets

There were 31 previously obtained hub targets of Sotetsuflavone for pancreatic cancer treatment all of which we analysed for KEGG enrichment in the DAVID database(https://david.ncifcrf.gov/summary.jsp). There were 119 signalling pathways enriched and the top 20 pathways with the lowest FDR values were selected for visualisation and analysis, and the enrichment of hub targets on each pathway was plotted in a Sankey diagram and a KEGG pathway enrichment bubble diagram (**Fig. 7**). The results of our study demonstrated that the major pathways of the hub targets associated with the treatment of pancreatic cancer with Sotetsuflavone are also closely related to the Pathways in cancer, Pancreatic cancer, and Prostate cancer. It is notable that both potential and hub targets were significantly enriched in pancreatic cancer signalling pathways. While further research is needed, this finding seems to indicated that Sotetsuflavone plays an important role in the treatment of pancreatic cancer.

**Fig. 7.**
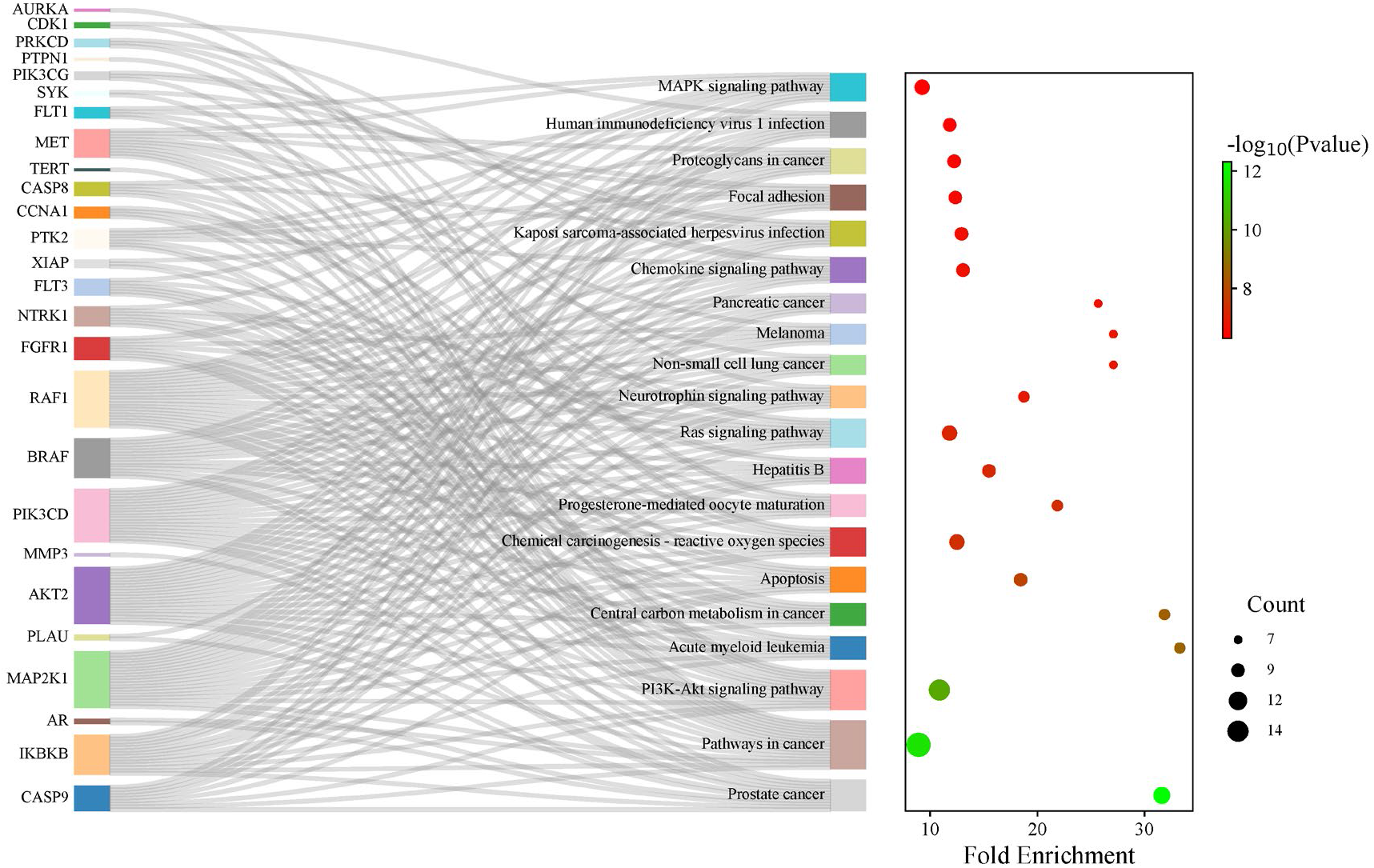
Enrichment of KEGG pathway for hub targets Sankey and bubble diagrams.

### 3.6. Survival analysis of hub targets

It is used by the Kaplan-Meier estimator in estimating the survival function. There is a Kaplan-Meier curve that can be used to show the probability of an event happening at a particular time[45].

The Kaplan-Meier survival curves were established and then log-rank of the survival rates were compared between the high and low expression groups using the log-rank test[19]. ABCB1, AURKA, CDK1, HDAC6, MET, and MMP3 were the 6 hub targets with p-value <0.01, which suggested that ABCB1, AURKA, CDK1, HDAC6, MET, and MMP3 might be involved in pancreatic survival, and further analysis of the RNA expression levels in normal and pancreatic cancer tissues was performed (**Fig. 8**).

**Fig. 8.**
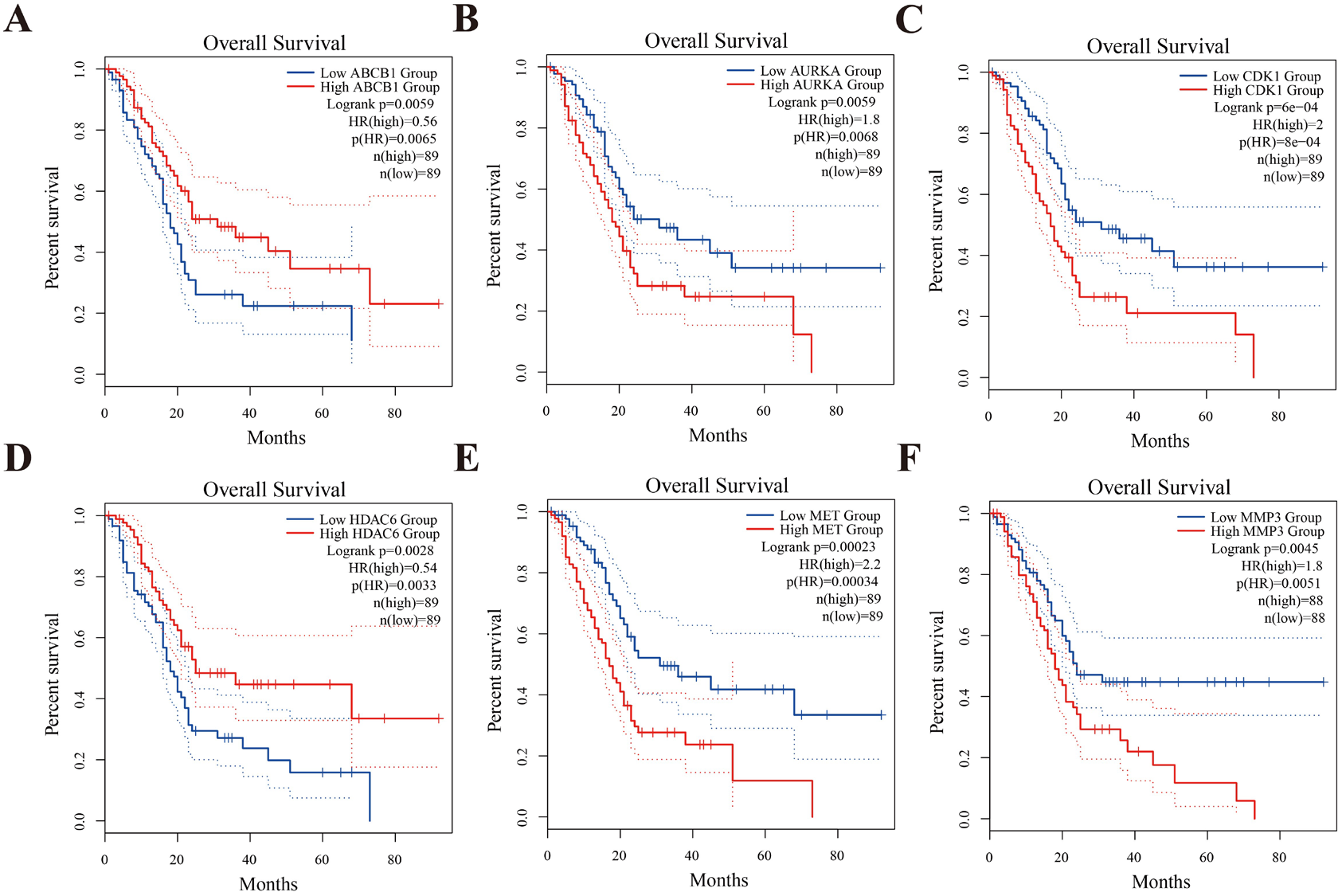
Using Pearson’s coefficient analysis at p < 0.01, the relationships between the six genes. (A)ABCB1, (B)AURKA, (C)CDK1, (D)HDAC6, (E)MET, (F)MMP3. Genes with no correlation are indicated in blue, genes with negative correlation are shown in red, and genes with positive correlation are shown in green.

### 3.7. Expression analysis of hub targets

The expression differences of ABCB1, AURKA, CDK1, HDAC6, MET and MMP3 at the RNA level in normal and pancreatic cancer tissues were analysed by the GEPIA 2 platform (http://gepia2.cancerpku.cn/).It was found that the RNA expression levels of ABCB1, AURKA, CDK1, HDAC6, MET and MMP3 were higher in tumour tissues than in normal tissues (**Fig. 9**). The above results suggest that high expression of ABCB1, AURKA, CDK1, HDAC6, MET and MMP3 in tissues is one of the causes of pancreatic cancer induction, but further studies are needed.

**Fig. 9.**
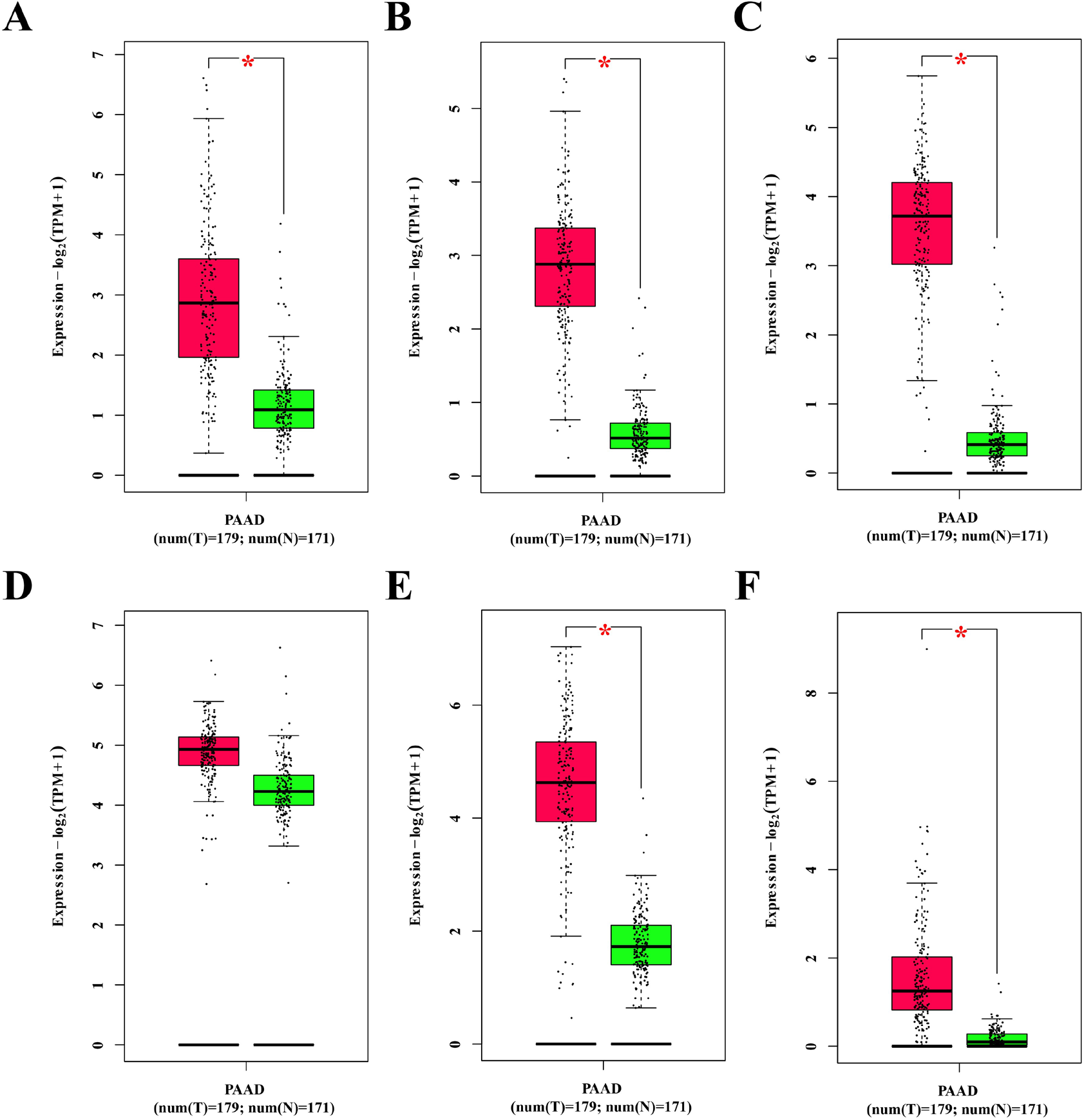
Expression analysis of hub targets in normal and PAAD tissues (p < 0.01). (A)ABCB1, (B)AURKA, (C)CDK1, (D)HDAC6, (E)MET, (F)MMP3.

### 3.8. Molecular docking of hub targets

With both survival and expression analyses, in which we selected 6 hub targets, ABCB1, AURKA, CDK1, HDAC6, MMP3, and MET, for molecular docking analyses. We downloaded protein crystal structures of 6 hub targets from the RCSB Protein Data Bank (PDB) (https://www.rcsb.org/) and used the ProteinsPlus platform (https://proteins.plus/) to predict the size and location of active pockets (**Table 2**). Firstly, the hub targets and Sotetsuflavone were pre-processed using AutoDock Tools 1.5.7. Then, molecular docking was done using AutoDock Vina and visual analyses were performed using Discovery studio visualizer, Maestro 12.8 and PyMOL.

**Table 2.** Molecular docking activity pockets information for targets.

| Targets | Uniport ID | Center X | Center Y | Center Z | Max. radius |
| --- | --- | --- | --- | --- | --- |
| ABCB1 | P08183 | 20.86 | 2.71 | -35.00 | 15.21 |
| AURKA | O14965 | -8.57 | 3.06 | 7.24 | 13.96 |
| CDK1 | P06493 | 6.77 | -6.11 | 2.3 | 21.24 |
| HDAC6 | Q9UBN7 | -1.38 | 1.23 | -12.71 | 51.43 |
| MET | P08581 | -5.57 | 1.34 | 26.33 | 19.29 |
| MMP3 | P08254 | -1.98 | 3.42 | -0.33 | 30.00 |

The interactions between Sotetsuflavone and 6 hub targets ABCB1, AURKA, CDK1, HDAC6, MET, and MMP3 were studied by molecular docking analysis. Molecular docking indicated that the binding energies of Sotetsuflavone to the 6 hub targets ABCB1, AURKA, CDK1, HDAC6, MET, and MMP3 were -8.6 kcal/mol, -8.8 kcal/mol, -4.2 kcal/mol, -10.4 kcal/mol, -10.7 kcal/mol and -8.0 kcal/mol, respectively (**Fig. 11**). As illustrated in **Fig. 10**, Sotetsuflavone interaction with Tyr310, Ile306, Phe303, Phe343, Phe336, Ala987 of the ABCB1 activity pocket. Sotetsuflavone interaction with Glu181, Glu260, Leu164, Phe144, Gly142, Lys162, Leu263, Val147, Leu210, Ala273 of the AURKA activity pocket. Sotetsuflavone interaction with Pro156, Arg127, Arg151 of the CDK1 activity pocket. Sotetsuflavone interaction with Asn783, Arg509, Ala1188, Thr1108, Pro1109, Val1107, Trp1182, Met1086, Gly1083 of the HDAC6 activity pocket. Sotetsuflavone interaction with Ala1108, Val1092, Met1211, Asp1222, Gly1087, Lys1110, Glu1127, Gln1123, Gly1224 of the MET activity pocket. Sotetsuflavone interaction with Pro59, Lys63, Pro373, Ser335, Asp43, His99, Glu66, Asp96 of the MMP3 activity pocket. The results showed that Sotetsuflavone had a strong binding ability to the 6 hub targets, and it was hypothesised that Sotetsuflavone could play a role in the treatment of pancreatic cancer by inhibiting the activities of the highly expressed hub targets and regulating the pancreatic cancer-related signalling pathways.

**Fig. 10.**
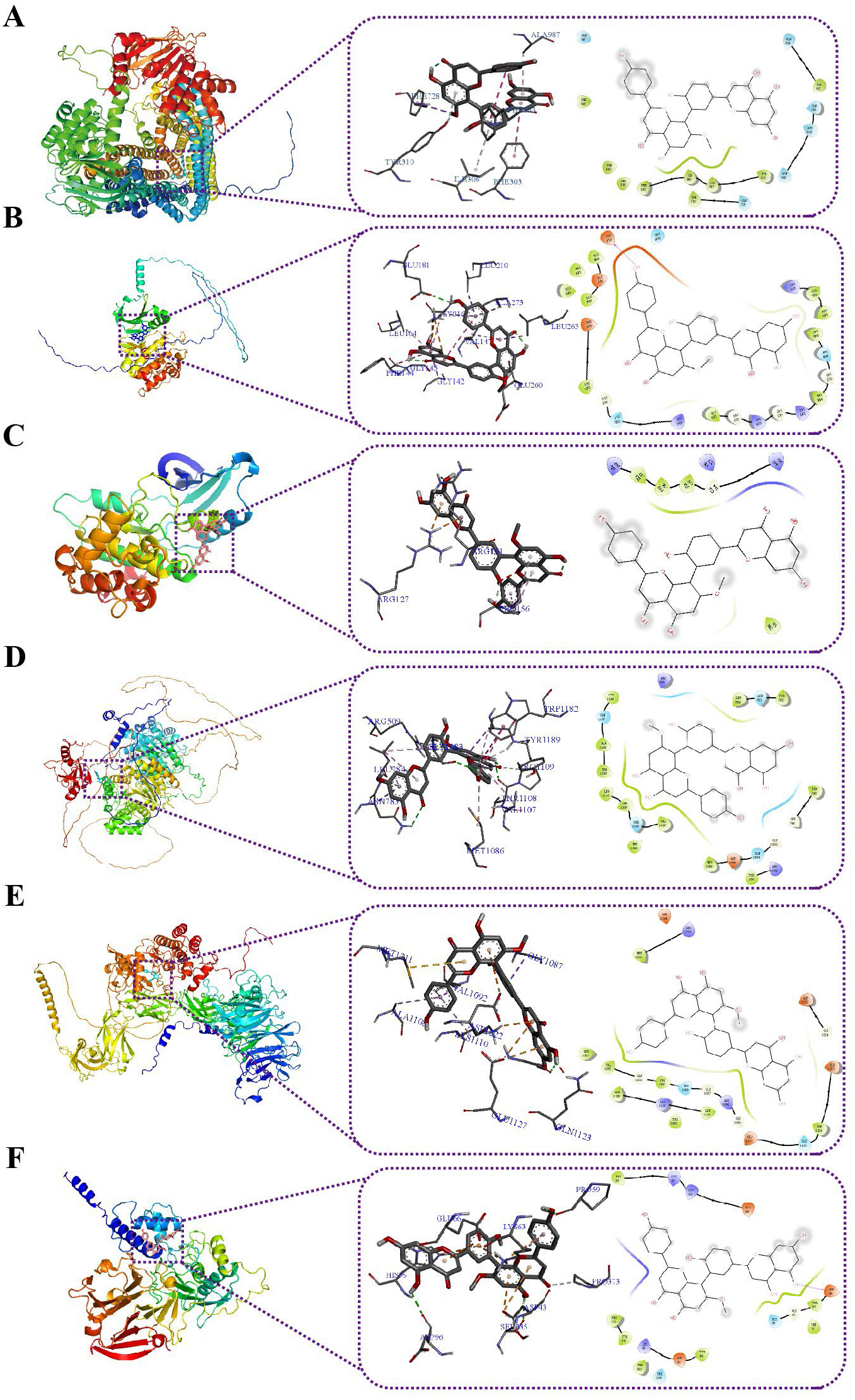
Visualisation of molecular docking results for hub targets. (A) Sotetsuflavone and ABCB1, (B) Sotetsuflavone and AURKA, (C) Sotetsuflavone and CDK1, (D) Sotetsuflavone and HDAC6, (E) Sotetsuflavone and MET, (F) Sotetsuflavone and MMP3.

**Fig. 11.**
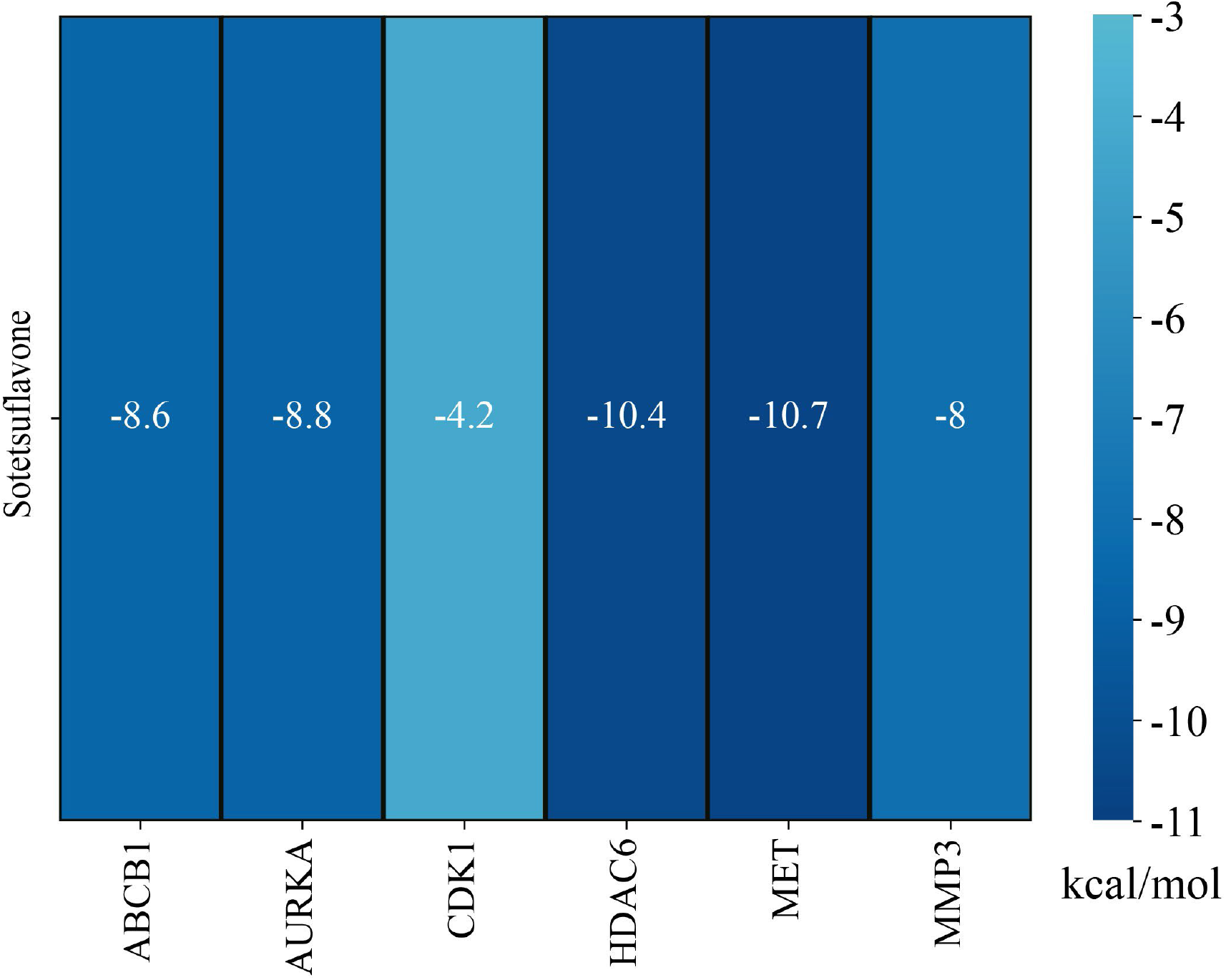
Heatmap of molecular docking binding energy of Sotetsuflavone to hub targets.

## 4. Discussion

There are studies which demonstrated that abnormalities in human microbial metabolism, blood type, blood glucose levels, lipid levels, gender, age, genetics, smoking, and alcohol consumption are all major contributing factors to the development of pancreatic cancer[2]. The use of surgical means is the primary way of treating pancreatic cancer[10]. Pancreatic cancer has limited treatment options and a high death rate, so the discovery of new anti-cancer drugs is urgent. In recent years, there has been widespread interest in the study of the pharmacological effects of natural products[46]. Network pharmacology contributes to the discovery of new drugs through helping us understand the complex mechanisms underlying the interactions of compounds with their targets[47]. In the present study, the first comprehensive pharmacological evaluation of Sotetsuflavone was carried out using multiple databases. In addition, we have also used network pharmacology, single-gene survival analysis, gene expression analyses and molecular docking to explore the mechanism of Sotetsuflavone in the treatment of pancreatic cancer, which lays the foundation for the research of Sotetsuflavone in the treatment of pancreatic cancer.

The comprehensive analysis of multiple databases discovered the characteristics of low toxicity, high bioactivity, good absorption and metabolism, and multiple targets of Sotetsuflavone (**Fig. 2**, **Fig. 3**). Sotetsuflavone has been reported to possess certain anticancer activities. Sotetsuflavone has been shown to induce autophagy in non-small cell lung cancer by blocking the PI3K/AKT/mTOR signalling pathway[13]. In addition, Sotetsuflavone could inhibit the invasion and metastasis of non-small cell lung cancer A549 cells by reversing EMT through TNF-α/NF-κB and PI3K/AKT signalling pathways[15]. Sotetsuflavone was also shown to inhibit proliferation and induce apoptosis in non-small cell lung cancer A549 cells through a ROS-mediated mitochondria-dependent pathway[11]. In addition, sotetsuflavone increased the inhibitory effect of 5-fluorouracil on gastric cancer cells through the AKT-mTOR pathway[16]. Therefore, Sotetsuflavone is promising to be a novel anticancer drug.

In the present study, we have identified Sotetsuflavone and its targets, which are involved in different cancer-related pathways, such as Prostate cancer, Pathways in cancer, and Pancreatic cancer (**Fig. 6**). It is worth noting that hub targets such as CDK1, MMP3, and MET are involved in these cancer-related signalling pathways. Network interactions analyses revealed that Sotetsuflavone can action on these hub targets, which indicated that Sotetsuflavone has anticancer activity (**Fig. 4**). It was concluded that the hub targets ABCB1, AURKA, CDK1, HDAC6, MET and MMP3 were involved in the survival of pancreatic cancer based on survival analyses (**Fig. 8**). In addition, it was found by gene expression analyses that the hub targets appeared to be highly expressed in pancreatic cancer tissues compared to normal tissues (**Fig. 9**). It is hypothesised that the high expression of these hub targets contributes to the induction of pancreatic cancer. It was further confirmed by molecular docking that Sotetsuflavone has a strong binding capacity to these hub targets. Therefore, it is assumed that Sotetsuflavone can inhibit the activity of these hub targets, and thus inhibit the signalling pathway related to pancreatic cancer to reach the therapeutic effect of pancreatic cancer.

In addition, GO enrichment analysis demonstrated that the anticancer targets of Sotetsuflavone were mainly associated with protein phosphorylation, negative regulation of apoptotic process, plasma membrane, receptor complex, protein kinase activity, and G-protein coupled receptor activity (**Fig. 5**). In addition, KEGG pathway enrichment analyses showed that the anti-cancer targets and hub targets of Sotetsuflavone were mainly involved in signalling pathways such as Prostate cancer, Pathways in cancer, PI3K-Akt signalling pathway, Central carbon metabolism in cancer, Apoptosis, Non-small cell lung cancer and Pancreatic cancer (**Fig. 6**, **Fig. 7**). It is reported in the papers that Sotetsuflavone can regulate the PI3K-AKT signalling pathway, Apoptosis and Non-small cell lung cancer signalling pathway to achieve anticancer effects[11, 13, 15]. The current study demonstrated that Sotetsuflavone can be used to treat pancreatic cancer by regulating pancreatic cancer and its associated signalling pathways.

In summary, there is scientific basis for our study to uncover Sotetsuflavone as a promising therapeutic option for pancreatic cancer. We have utilised network pharmacology to demonstrate the multi-target pharmacological mechanisms of Sotetsuflavone in pancreatic cancer. Furthermore, we utilised survival analysis, gene expression analysis and molecular docking analysis to validate our results. However, there is a necessity for more in vivo and in vitro studies to validate the validity of the present discoveries so that the mechanisms of Sotetsuflavone in the treatment of pancreatic cancer can be more clearly elucidated.

## 5. Conclusions

During the recent decades, pancreatic cancer has been on the increase around the world. Other than surgical treatment, there has not been much improvement. In the present study, we have used multiple databases to perform a comprehensive pharmacological evaluation of Sotetsuflavone and identified several novel targets for its treatment of pancreatic cancer. The mechanism of Sotetsuflavone treatment for pancreatic cancer was revealed using network pharmacology. This mechanism was validated through further single gene survival analysis, gene expression analysis and molecular docking analysis. The results showed that the binding ability of Sotetsuflavone to the hub targets ABCB1, AURKA, CDK1, HDAC6, MET, and MMP3 was strong, and the hub targets were involved in pancreatic cancer-related signalling pathways, which suggests that the hub targets ABCB1, AURKA, CDK1, HDAC6, MET, and MMP3 are the potential targets of Sotetsuflavone for the treatment of pancreatic cancer. In conclusion, Sotetsuflavone can play its therapeutic role in pancreatic cancer through multiple pathways acting on multiple targets. The present study establishes a framework for further research on the protective mechanism of Sotetsuflavone in the treatment of pancreatic cancer, as well as a foundation for the development of novel anticancer drugs.

## CRediT authorship contribution statement

Zi-Yong Chu designed the experiment and performed the statistical analysis. Zi-Yong Chu and Xue-Jiao Zi carried out the study. Zi-Yong Chu and Xue-Jiao Zi drafted the manuscript. All authors read and approved the final manuscript.

## Acknowledgement

The authors declare that they have no known competing financial interests or personal relationships that could have appeared to influence the work reported in this paper. This research did not receive any specific grant from funding agencies in the public, commercial, or not-for-profit sectors.

## Data availability

The data that has been used is confidential.

